# The analysis of a large dataset of PCR-test results in the community reveals the impact of age, variants and vaccination status on SARS-CoV-2 viral dynamics

**DOI:** 10.64898/2025.12.04.692282

**Authors:** Maxime Beaulieu, Nathanaël Hozé, Vincent Vieillefond, Thimothée Goetschy, Gina Cosentino, François Blanquart, Florence Débarre, Jérémie Guedj

## Abstract

The COVID-19 pandemic has shown the value of large-scale community PCR tests for epidemic surveillance, but the viral load measurements they provide have seldom been exploited to reconstruct within-host trajectories. Because these data are collected for diagnostic rather than research purposes, they are characterized by sparse longitudinal follow-up, heterogeneous sampling, and missing metadata, raising uncertainty about their usefulness to reconstruct with-host viral dynamics.

We first conducted a simulation study to assess the feasibility of estimating the viral dynamics patterns from such datasets. Across multiple scenarios replicating realistic sampling patterns, we found that peak viral load and clearance time could be estimated with good accuracy, although uncertainty tended to be underestimated. Parameters driving early viral kinetics, e.g. incubation and proliferation time, were mostly estimated with poor precision, as most community tests are conducted after symptom onset, later in infection.

We then applied this framework to a large dataset of 322,218 PCR tests associated with symptomatic SARS-CoV-2 infections in France between July 2021 and March 2022, encompassing both Delta-variant circulation and the emergence of first Omicron variants. We quantified the effects of age, vaccination, and variants on viral load trajectories. Age ≥65 years was consistently associated with longer clearance time, extending the duration of detectable viral load by 2 to 6 days. Vaccination shortened the clearance time by 2 to 4 days but had minimal impact on peak viral load. Infections by Omicron-variants were associated with a lower peak viral load and shorter clearance times compared with pre-Omicron (Delta) infections.

Thus, community PCR tests can be leveraged to identify key parameters of viral dynamics. As multiplex PCR testing becomes increasingly widespread, establishing robust frameworks for data collection, sharing, and privacy protection will be essential to support the use of these data.

**Author Summary:** In this study, we explored whether viral load data collected routinely in community laboratories can help us understand how factors like age, vaccination, and viral variant influence the course of infection. These data, collected for diagnostic purposes rather than research, are large but often incomplete, with most individuals tested only after the time of symptoms onset. We first used simulations to assess whether, despite these limitations, key aspects of viral dynamics can be reliably estimated. We then applied our approach to millions of test results collected in France during periods dominated by Delta and Omicron variants. We found that older age was consistently associated with longer infection duration, while vaccination reduced the time the virus remained detectable without affecting peak viral levels. Omicron infections were generally associated with lower peak viral load and somewhat faster clearance than pre-Omicron infections. Overall, our work demonstrates that community testing data, can provide valuable insights into viral dynamics and help monitor the effects of new variants or interventions. As multiplex tests become more common, facilitating and improving data collection, sharing, and privacy protection will be essential to make the most of these resources in future outbreaks.

## Introduction

Virological tests done in the general community have become an essential component of the public health surveillance system during an outbreak. During the Covid-19 pandemic, the design, volume and level of detail of these data have also opened new avenues for mathematical modellers. As a striking example, theoretical modelling performed in 2021 showed that the analysis of random large scale cross sectional distribution of SARS-CoV-2 viral loads could inform on the dynamics of the epidemics(1).

The emergence of variants of concern (VOCs) and the implementation of large-scale vaccination campaigns have largely influenced both the severity and the transmission of SARS-CoV-2 (2–5). Yet, and despite dozens of studies, the impact of Omicron emergence and vaccination on viral load remains debated. In general, there is a consensus towards an effect of vaccination on viral load with pre-Omicron variants (6–8). The effect of vaccination on Omicron variants is less clear, with some studies suggesting that vaccination may reduce viral load (7) while other suggest no effects (8–10).

Here we hypothesize that the millions of PCR tests done in the general population could be analysed in a systematic and quantitative way to leverage the information and reduce the potential bias present in small or retrospective cohort studies. Identifying signals associated with a change in viral dynamics is nonetheless highly challenging with such data. First, these data have been collected for diagnostic, and not for research purposes. Therefore, they are typically characterized by a large proportion of missing metadata, high variability in sample collection and laboratory procedures, making it challenging to compare and analyse viral load values. In addition, sampling times are skewed, with most individuals being sampled close to symptom onset, and only very few longitudinal follow-ups, questioning whether such data can be used to reliably infer on viral dynamics.

In this study, we first evaluated through simulation the feasibility of inferring viral dynamics from sparse and heterogeneous data. We then analysed a large dataset of community-based PCR tests collected in France in symptomatic individuals during the circulation of both Delta and Omicron variants to assess the impact of variant, vaccination status, and age on viral load dynamics.

## Materials and methods

### Data collection

The dataset includes all PCR tests performed in Biogroup laboratories, one of the largest groups of private community laboratories in France, between July 1, 2021, and March 13, 2022 (**Figure S1**). This dataset extends a previously described dataset by integrating additional data collected during summer and autumn 2021(11–13). Only individuals for which the following information were available were analysed: cycle threshold (Ct) value, sex, age, vaccination status, symptomatic status, time since symptom onset (see below). All these pieces of information (except Ct value) were self-reported.

As these data were not collected for research purposes, we made a number of assumptions to make them suitable for analysis. We considered an individual as infected if at least one positive PCR test with quantified Ct value was available. All tests with the same variant of infection within 30 days after the first detection were considered to be the same infection. When available, viral sequencing was used to determine the variant of infection; otherwise, we relied on mutation information obtained from the PCR tests (**Table S1-S2**). When the virus genotype was not available, the variant of infection was assumed to be the dominant variant in the population, i.e., representing more than 95% of circulating variants at the time of symptom onset (14,15). As Beta and Gamma, as well as Omicron BA.1 and BA.2, cannot be distinguished by mutation profiles, we classified variants only as pre-Omicron or Omicron. Notably, the vast majority of pre-Omicron infections corresponded to the Delta variant. The time of symptom onset was reported by intervals (0–1, 2–4, 5–7, 8–14, 15–28, and >28 days) and we used in the model the middle of the interval as the time of onset. Individuals were considered vaccinated upon self-report of receiving at least one dose of the vaccine.

### Modelling viral kinetics

We characterized the viral kinetics, from time of infection to viral clearance using a piecewise linear mixed-effects model, which we fitted to cycle thresholds (**Figure 1**). The time of infection is reconstructed from the incubation period (*T*_*I*_), estimated from the time of symptom onset. Upon infection, the viral load increases for a duration T_P_, called the proliferation period, and the viral load reaches its peak noted V_P._ Then the viral load declines to clearance for a duration noted T_C_. Let ***Y***_***i***_ = {(*y*_*i*,1_,*t*_*i*,1_), …, (*y*_*i*,*j*_, *t*_*i*,*j*_)} denote the vector of observations for the individual *i*, with the *j*^*t*ℎ^ Ct value *y*_*i*,*j*_ observed at time *t*_*i*,*j*_ since symptom onset, described by the following equations:

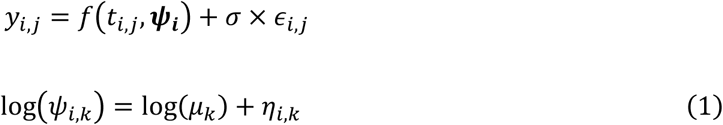

**Figure 1:**
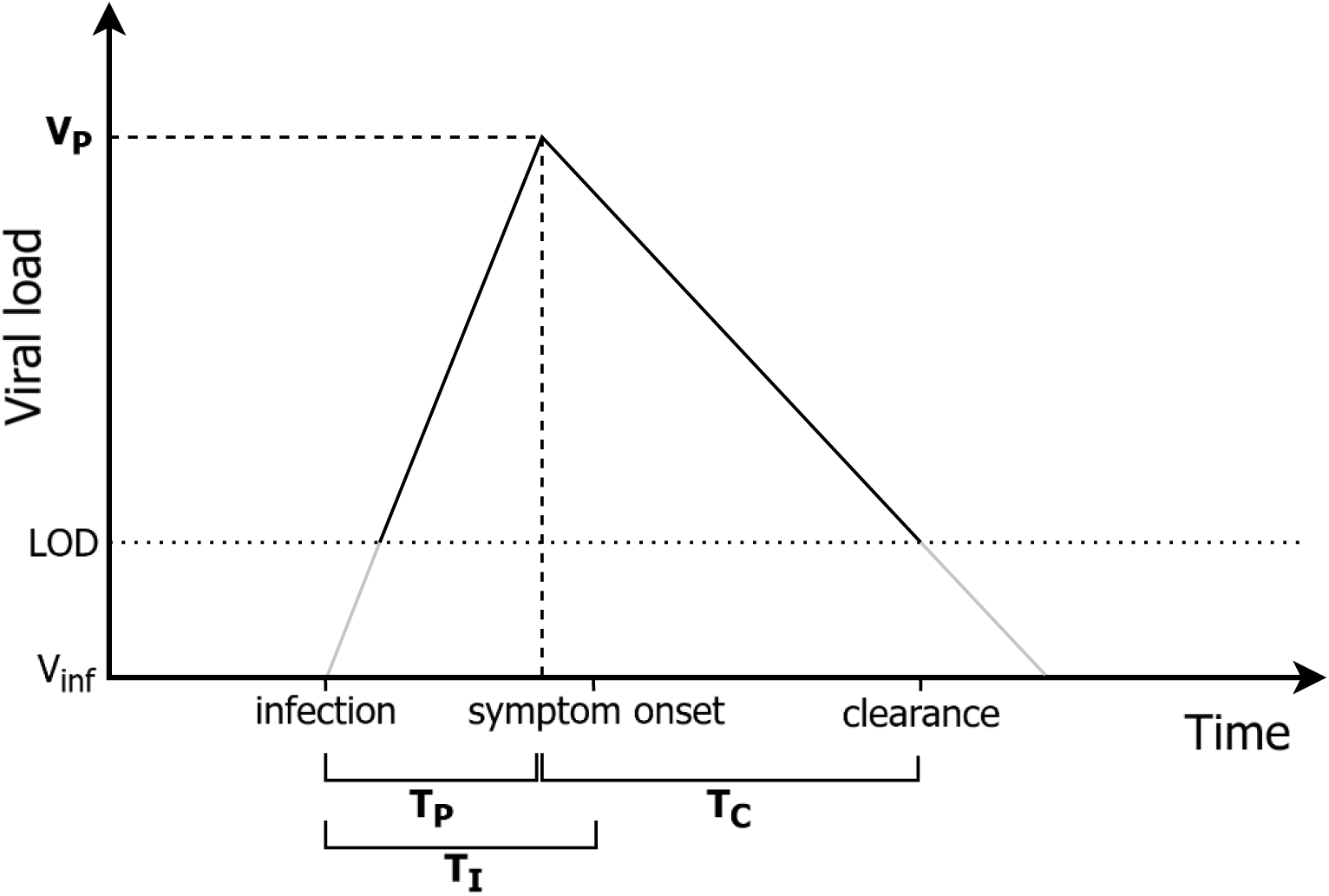
Schematic of the piecewise linear viral dynamics model. Schematic representation of viral kinetics from infection to clearance, showing in bold the estimated parameters: incubation period (T_I_), proliferation period (T_P_), peak viral load (V_P_), and clearance period (T_C_). LOD denotes the limit of detection, and V_inf_ the viral load at the time of infection.

where *f*(*t*_*i*,*j*_, ***ψ***_***i***_) represents the structural model, a function of *t*_*i,j*_, and ***ψ***_***i***_ = {*T*_*I_i_*_, *T*_*P_i_*_, *V*_*P_i_*_, *T*_*C_i_*_} the vector of individual parameters. Each individual parameter *ψ*_*i*,*k*_ (with *k* ∈ {1,.., 4} the parameter index) is composed of *μ*_*k*_, a fixed effect (or population parameter) shared by all the individuals, and *η*_*i*,*k*_, the random effect of the parameter *k* specific to the individual *i*, which is assumed to follow a lognormal distribution with variance-covariance matrix Ω. We considered a residual error model with σ the standard deviation of the residual error, and *∈*_*i,j*_∼*N*(0,1). We set the Ct value at time of infection to 50, which is 10 Ct above the limit of detection.

### Simulation study

#### Scenarios

We conducted a simulation study to evaluate the performance of parameter estimation in terms of bias, uncertainty, and computational efficiency. A total of 50 datasets were simulated with various population sizes, but keeping constant the ratio of 50% of individuals being infected and 50% being non-infected. The viral kinetics of infected individuals were simulated using the same population parameters: an incubation period (*μ*_*T_I_*_) of 5 days, a proliferation period (*μ*_*T_P_*_) of 6 days, a peak viral load (*μ*_*V_P_*_) of 25 Ct, and a clearance period (*μ*_*T_C_*_) of 15 days. The standard deviation of individual random effects (*η*_*i*,*k*_) was set to 0.15 for all parameters. The measurements were assumed to be noisy, with a variance of 4 Ct, of the same order of magnitude as that estimated from the real dataset (**Figure S2**). The sensitivity of PCR tests is imperfect and decreases as viral load decreases. Thus, modelling all individuals regardless of whether they have a positive PCR test enables the screening of infected individuals who only test negative because they were sampled beyond the window of detectable viral load.

Three scenarios for the sampling times were considered. In the first scenario, we assumed a rich sampling design in a population of 120 individuals (i.e. 60 infected individuals), with daily observations from infection to clearance for each individual. Including only individuals with at least one positive test guarantees that all infections are captured, since viral load is expected to be PCR-detectable for at least one day during the infection. The second scenario assumed a population of 4,000 individuals (i.e. 2,000 infected individuals) and that the sampling times were uniformly distributed during the infection period, i.e., from one week before symptom onset to three weeks after symptom onset. The distribution of the number of tests per individual was similar to that in the real dataset (1 test: 84%, 2 tests: 14%, 3 tests: 2%, 4 and more: <1%), reflecting limited longitudinal measurements. Restricting the analysis to individuals with at least one positive test captures only 80–85% of all infections. The third scenario assumed a population of 4,000 individuals (i.e. 2,000 infected individuals), and best reflected the data collected in community laboratories, with the distribution of the number of tests and sampling times both similar to those in the real dataset. Thus, in this scenario tests were mainly performed just after the onset of symptoms (see **Figure 4**). Restricting the analysis to individuals with at least one positive test captures more than 99% of the infections.

#### Inference

In all three scenarios, we evaluated two different modelling approaches. The first approach consisted in modelling only individuals with at least one positive PCR test, assuming that they were all genuinely infected. Thus, we excluded from the modelling all individuals with exclusively negative PCR tests. With this approach, the likelihood of observing ***Y***_***i***_ can be written as follows:

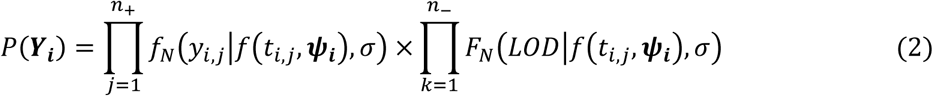

where *f*(*t*_*i*,*j*_, ***ψ***_***i***_) is the predicted Ct value under the viral dynamics model, *f*_*N*_(*y*_*i*,*j*_|*f*(*t*_*i*,*j*_, ***ψ***_***i***_), *σ*) the density of the Normal distribution evaluated at the *j*^th^ observation of individual *i*, *y*_*i*,*j*_, with mean *f*(*t*_*i*,*j*_, ***ψ***_***i***_) and observation error *σ*, *F*_*N*_(*LOD*|*f*(*t*_*i*,*j*_, ***ψ***_***i***_),*σ*) the Normal cumulative distribution function evaluated at the limit of detection (LOD)(16).

The second approach consisted in modelling both individuals with at least one positive PCR test, and individuals with only negative tests, thereby including individuals who may not be infected. In this situation, both the infection status of each individual and the viral kinetic parameters of infected individuals are inferred. The infection status of each individual can be inferred by integrating PCR test results with their timing relative to symptom onset. In this case the likelihood of observing ***Y***_***i***_ is expressed as the sum of contributions from the cases where the individual is infected vs. not infected:

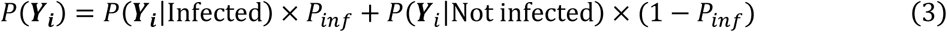

where *P*(***Y***_***i***_|Infected) is defined in (**Eq2**), *P*_*inf*_ denotes the estimated proportion of infected individuals, and *P*(***Y***_*i*_|Not infected) is defined as follows:

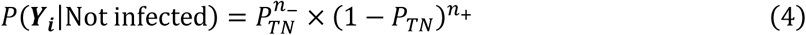

Here, *n*_+_ and *n*_−_ are the number of positive and negative PCR tests for each individual, and *P*_*TN*_ the probability of true negative, fixed to 0.9998, in line with the high specificity of PCR testing(17).

We estimated the model parameters under two different inference frameworks: a Bayesian framework and a frequentist framework. The Bayesian inference relied on the HMC-NUTS algorithm(18) implemented in Stan (via the rstan package, version 2.32.6(19)), and was applied to both modelling approaches. The frequentist framework used the stochastic approximation expectation– maximization (SAEM) algorithm(20) implemented in Monolix 2023R1(21), but could only be applied to the first approach (i.e., restricted to infected individuals), because the likelihood accounting for infection status cannot be specified in Monolix. The simulation codes and models are available in the repository (22).

In Stan, we ran four chains in parallel, with random initial value sampled in the priors, with 400 iterations each including 200 warm-up iterations, leading to a posterior sample of 800 replicates. We considered the mean of the posterior distribution as the estimate of the parameter. We used moderately informative priors, using Gaussian distributions centred on the simulated values (**Figure S3**). We restricted the analysis to fits with convergent chains, defined by the criterion R^ < 1.05(23), which compares the between- and within-chain estimates.

#### Evaluation

The performance of both frequentist and Bayesian inference, for each modelling approach in each scenario, was evaluated by the relative error of estimation (REE), the coverage rate of the 95% confidence (or credibility) intervals of the parameters, and computation time (24).

The relative error of estimation assesses the accuracy of estimation, and is defined as:

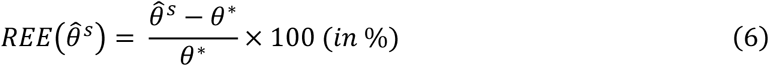

With 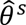 denoting the estimate of parameter *θ* (posterior mean for Bayesian inference, or maximum likelihood estimate for frequentist inference) in simulation *s*, and *θ*^∗^ the true simulated value of *θ*.

Coverage rates were used to evaluate the validity of the 95% confidence (or credibility) intervals, reflecting both the potential bias and the uncertainty of parameter estimates. The coverage rate was calculated from 50 simulations in which the population parameters were identical across all scenarios and datasets. The 95% confidence interval of the coverage rate was computed assuming a binomial distribution with number of trials 50. The nominal target coverage is 0.95, and intervals including this value suggest appropriate uncertainty quantification.

### Inference in PCR data in the general community

Only symptomatic individuals with at least one positive PCR test were included in the model, with all data related to their infection (see above). All observations below the LOD were considered as censored. The analysis focused on assessing the influence of variant of infection (Pre-Omicron vs. Omicron), vaccination status (vaccinated vs. unvaccinated), and age (<65 vs. ≥65 years) on viral load dynamics. Given the large number of individuals (**Table 1**), and the potential interaction between these covariates, a composite covariate with eight subgroups, defined by the combinations of these three factors, was constructed. Thus, a single parameter inference was performed, and the population parameters (the fixed effects *μ*, the variance of random effects *η*, and the residual error *σ*) were shared by subgroups. Covariates effects on viral dynamics were illustrated using a forest plot of parameter estimates with their 95% confidence intervals for each subgroup, together with the predicted mean trajectory and its 95% confidence interval. Due to the prohibitive computational time for a dataset of this size (**Figure S6**), parameter estimation was performed using the stochastic approximation expectation-maximization (SAEM) algorithm implemented in Monolix 2023R1.

**Table 1:** Description of the study population.

| Age | < 65 years old |  |  |  | ≥ 65 years old |  |  |  |
| --- | --- | --- | --- | --- | --- | --- | --- | --- |
| Variant of infection | Pre-Omicron |  | Omicron |  | Pre-Omicron |  | Omicron |  |
| Vaccination status | Not vaccinated | Vaccinated | Not vaccinated | Vaccinated | Not vaccinated | Vaccinated | Not vaccinated | Vaccinated |
| Number of individuals | 44,762 | 35,875 | 53,349 | 166,867 | 2,022 | 4,349 | 1,613 | 13,381 |
| Female | 23,520 | 20,249 | 28,167 | 96,333 | 1,256 | 2,449 | 987 | 7,553 |
| Number of tests per individual |  |  |  |  |  |  |  |  |
| 1 | 39,289 | 31,055 | 44,862 | 137,333 | 1,725 | 3,725 | 1,302 | 10,119 |
| 2 | 4,787 | 4,212 | 7,532 | 25,188 | 246 | 520 | 264 | 2,570 |
| 3 | 601 | 541 | 849 | 3,663 | 37 | 97 | 40 | 562 |
| ≥ 4 | 85 | 67 | 106 | 683 | 14 | 7 | 7 | 130 |
| Time since symptom onset, per test (days) |  |  |  |  |  |  |  |  |
| [0-1] | 12,393 | 9,541 | 22,398 | 53,289 | 443 | 861 | 464 | 3,482 |
| [2-4] | 25,116 | 21,444 | 25,762 | 97,415 | 1,100 | 2,549 | 848 | 8,284 |
| [5-7] | 5,451 | 3,787 | 4,259 | 13,552 | 330 | 651 | 222 | 1,217 |
| [8-14] | 1,455 | 891 | 769 | 2,185 | 120 | 233 | 70 | 324 |
| [15-28] | 242 | 158 | 102 | 296 | 22 | 41 | 6 | 51 |
| ≥ 28 | 105 | 54 | 59 | 130 | 7 | 14 | 3 | 23 |

## Results

### Simulation study

The first simulation scenario considered a rich sampling design (daily Ct values) in a cohort of N=120 individuals (50% being infected), and, as expected in such a rich setting, all parameters were estimated with good precision and accuracy and coverage of 95% in both Bayesian and frequentist inference frameworks (**Figure 2A and 3A**).

**Figure 2:**
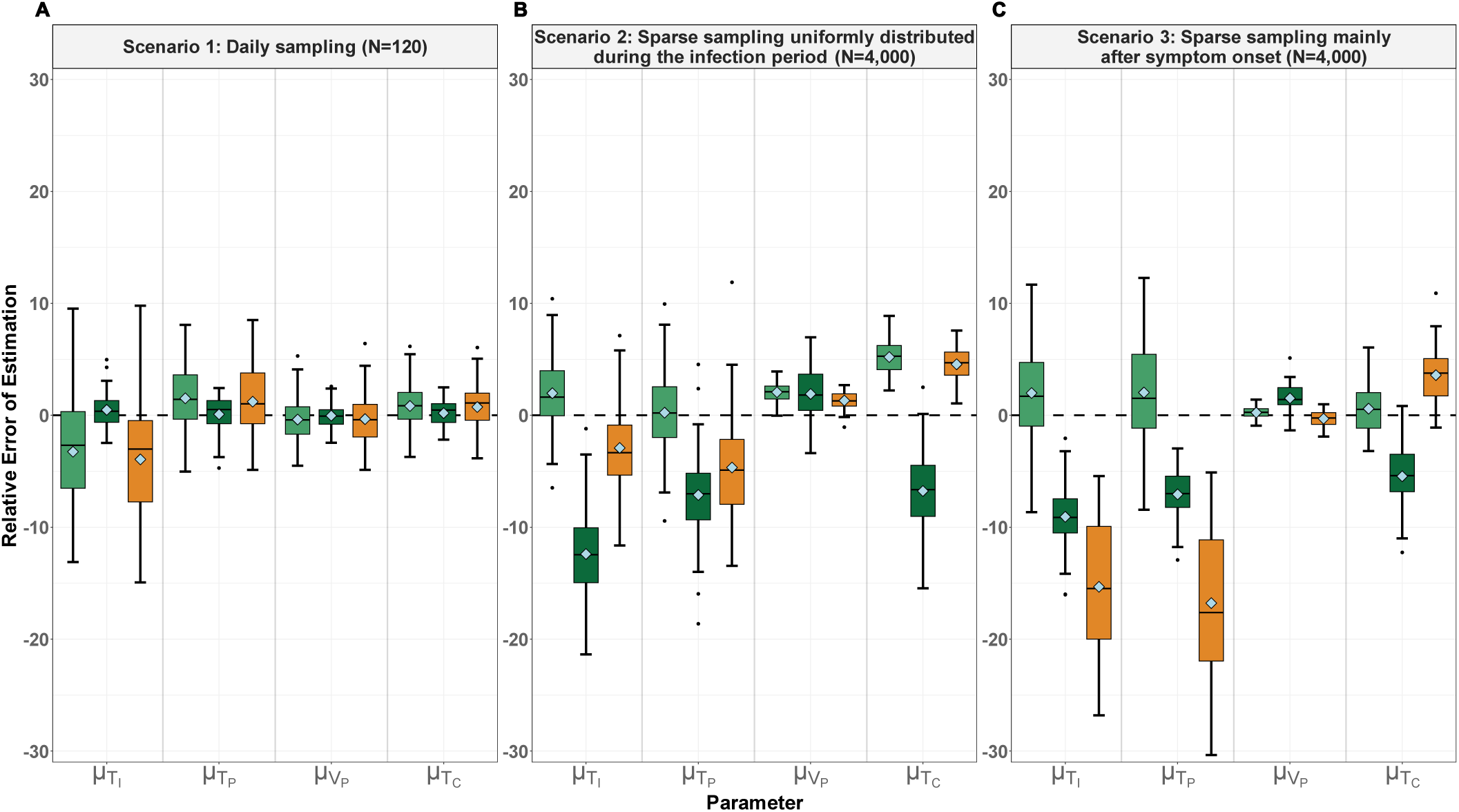
Estimation of viral dynamics parameters from simulation studies. Boxplots represent the distribution of the relative error of estimation of the population parameters for the incubation period (*μ*_*T_i_*_, in days), the proliferation period (*μ*_*T_P_*_, in days), the peak viral load (*μ*_*V_P_*_, in Ct), and the clearance period (*μ*_*T_C_*_, in days). The central line in each box indicates the median, the box represents the interquartile range (IQR; 25%–75%), and the whiskers extend to 1.5 × IQR or to the most extreme values. Values beyond this range are shown as outliers (black dots), while the diamond represents the mean of relative error of estimation (relative bias). Results are given for three scenarios of data generation (see methods). Scenario 1: daily observations from infection to clearance in 120 individuals (50% infected); scenario 2: sparse data uniformly distributed from infection to clearance in 4,000 individuals (50% infected); scenario 3: sparse data mainly after symptom onset in 4,000 individuals (50% infected). Light green: both positive and negative tests for all individuals are included in the Bayesian framework using Stan; dark green: only positive individuals are included in the Bayesian framework using Stan; orange: only positive individuals are included in the frequentist framework using Monolix.

**Figure 3:**
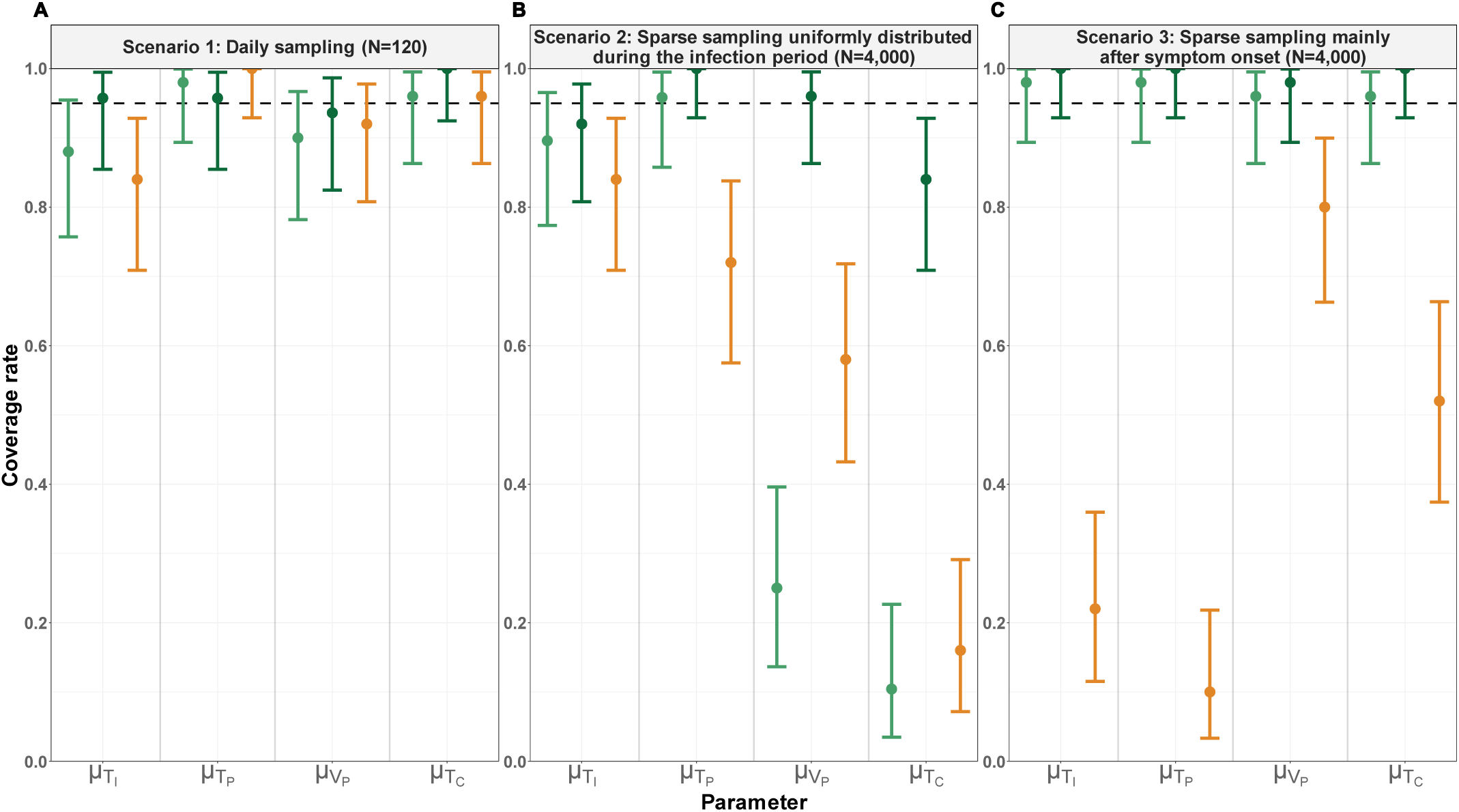
Coverage rates of the viral dynamics parameters. The dots (bars) represent the coverage rate (and 95% confidence interval) of the population parameters for the incubation period (*μ*_*T_i_*_, in days), the proliferation period (*μ*_*T_P_*_, in days), the peak viral load (*μ*_*V_P_*_, in Ct), and the clearance period (*μ*_*T_C_*_, in days). Scenario 1: daily observations from infection to clearance in 120 individuals (50% infected); scenario 2: sparse data uniformly distributed from infection to clearance in 4,000 individuals (50% infected); scenario 3: sparse data mainly after symptom onset in 4,000 individuals (50% infected). Light green: both positive and negative tests for all individuals are included in the Bayesian framework using Stan; dark green: only positive individuals are included in the Bayesian framework using Stan; orange: only positive individuals are included in the frequentist framework using Monolix.

We then tested a more realistic scenario with N=4,000 individuals (50% being infected), with most individuals having only one observation, randomly sampled during the infection period (see methods). The mean relative error was larger than in the first scenario but remained below 15% for all parameters, regardless of the modelling approach and inference framework, and equal to 2 and 5% for peak viral load (*μ*_*V_P_*_) and time to clearance (*μ*_*T_C_*_), respectively, corresponding to an absolute error of 0.5 Ct and 0.8 day, respectively (**Figure S4B**). Interestingly, the coverage rates were lower than 95% for most parameters, regardless of the modelling approach and inference framework, with the lowest values equal to 25 and 10% for peak viral load and time to clearance, respectively, (**Figure 3B**). Together these findings indicate that in presence of a large population of individuals with random sampling times, all methods tended to underestimate uncertainty, but the parameters of viral kinetics could be accurately estimated, with low absolute errors (**Figure S4B**).

Finally, the third scenario mimicked the real dataset with N=4,000 individuals (50% being infected), with most individuals having only one observation, sampled mainly after symptom onset. In spite of the skewed distribution in sampling times, the mean relative error was close to that observed in the second scenario, with values below 20% for all parameters. Errors were larger for the approach restricted to individuals with at least one positive PCR test. The mean relative error did not exceed 2 and 5% for peak viral load (*μ*_*V_P_*_) and time to clearance (*μ*_*T_C_*_), respectively, corresponding to an absolute error of 0.4 Ct and 0.8 day, respectively (**Figure S4C**). The coverage rates were improved for peak viral load and time to clearance, equal to 80 and 52%, respectively (**Figure 3C**), thanks to denser sampling post-symptoms. Conversely, as almost no information was available before symptom onset, the coverage rate was largely deteriorated for parameters driving the early viral kinetics, with values equal to 22 and 10% for the incubation period (*μ*_*T_i_*_) and the proliferation period (*μ*_*T_P_*_), respectively, using a frequentist framework. The coverage rate was much better when using a Bayesian framework, reflecting the strong influence of priors in this setting. Detailed results on the relative estimation errors for all parameters across the scenarios are presented in the Supplementary Material (**Figure S5**).

Finally, while the Bayesian inference framework with Stan provided a better estimation of the uncertainty, it came at a substantial computational cost due to the large sample size and the use of the HMC-NUTS algorithm. In Stan, when modelling the 2,000 individuals with confirmed infection, runtimes were approximately 40 times longer than those in Monolix (3.5 h vs 5.4 mins; **Figure S6**). The analysis of the full population of 4,000 individuals was extremely cumbersome, with an average runtime of 14 hours in the third scenario, limiting the use of Bayesian inference for our data. Consequently, we performed parameter estimation on the large community-based PCR dataset using the SAEM algorithm implemented in Monolix 2023R1, which provided reliable estimates of key parameters (peak viral load and clearance period) with minimal bias and acceptable computation times.

### Inference of the within-host kinetics of infections by pre-Omicron and Omicron variants in the general community

#### Description of the population

We then applied our framework to a large real-life dataset of all PCR tests results done in Biogroup community laboratories in France between July 1, 2021, and March 13, 2022. The initial dataset comprised 6,668,557 PCR tests from 4,242,747 individuals—a remarkably large dataset thanks to the very frequent testing in that period in France. After data processing and cleaning (see methods), 383,874 positive tests were analysed, corresponding to 322,218 symptomatic infections, for which age, sex, variant of infection, vaccination status, and time since symptom onset were available (**Figure S7**). The Pre-Omicron infection subgroups consisted predominantly of Delta infections, which was by far the dominant circulating variant in France during the summer and autumn of 2021 (14,15).

The largest group involved vaccinated Omicron cases aged <65 years (N = 166,867), followed by unvaccinated Omicron cases aged <65 years (N = 53,349), while the smallest groups were unvaccinated individuals aged ≥65 years with pre-Omicron (N = 2,022) or Omicron (N = 1,613 for Omicron) infections. Most positive tests were within 4 days after symptom onset (**Figure 4**), and only a small fraction of individuals, 16%, had repeated measurements (**Table 1**). In the population of individuals with at least one positive test, the virus cleared progressively, with 25 and 62% of individuals turning negative during the second week and the third week after symptom onset respectively (**Figure 4**).

**Figure 4:**
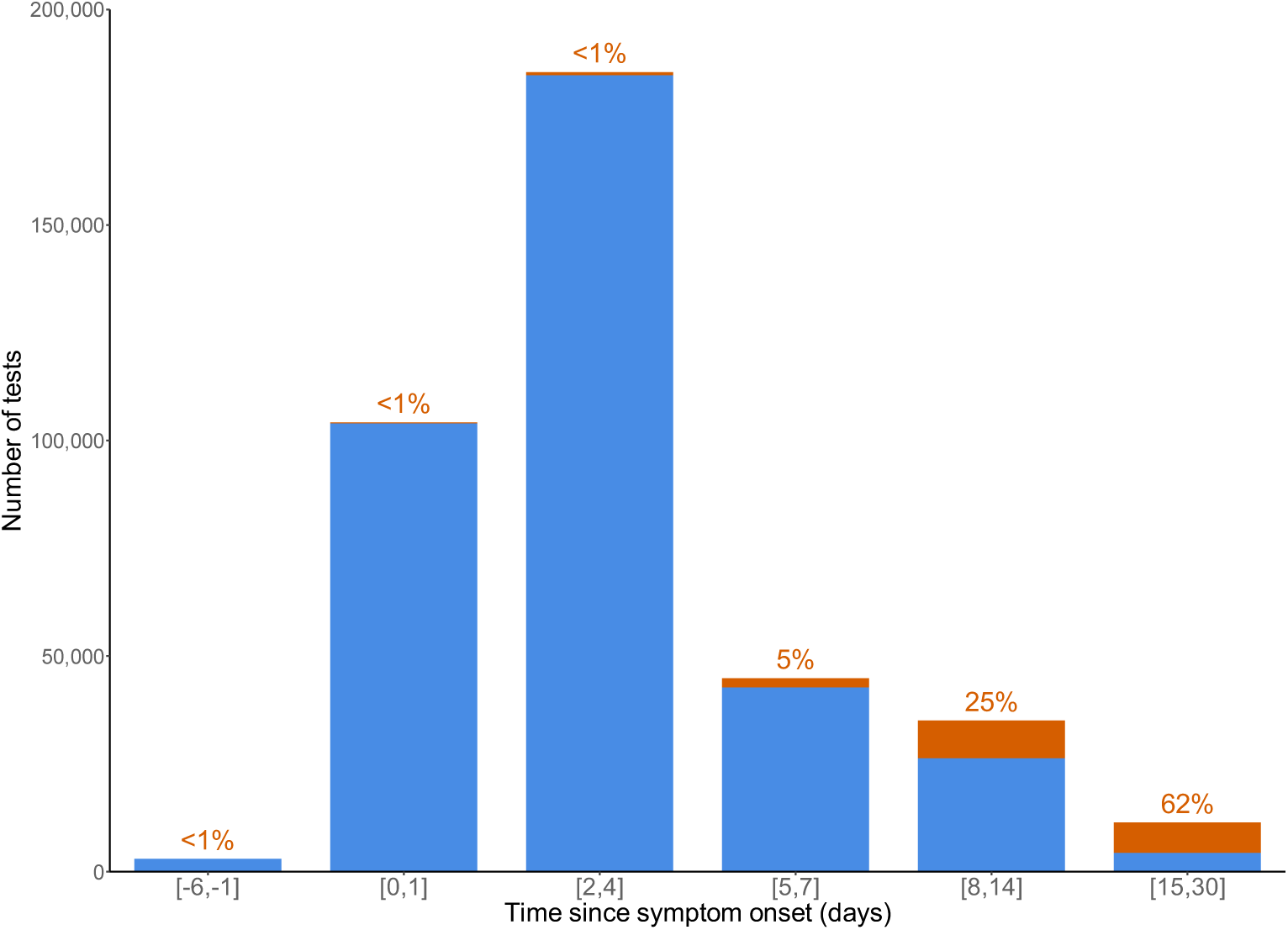
Distribution of negative and positive PCR tests in infected individuals. Number of positive and negative PCR tests available according to the time since symptom onset, and percentage of negative PCR tests. Blue: positive PCR tests; orange: negative PCR tests.

#### Estimation of the viral load dynamics parameters

To analyse the large community dataset, we estimated model parameters using the maximumlikelihood estimator obtained with the SAEM algorithm implemented in Monolix Software 2023R1. Stan was not used because computation times were prohibitively long for a dataset of this size (**Figure S6**), and Monolix yielded estimates of the key parameters (peak viral load and clearance period) with minimal bias (**Figure 2C**).

The model fitted distinct Ct trajectories across the eight subgroups (**Table S3**). Estimation uncertainty was slightly higher in older age classes, particularly for incubation and proliferation phases, reflecting the smaller sample sizes in these groups.

The clearance period was consistently increased across variants and vaccination groups for individuals older than 65, by approximately 2 to 5 days (**Figure 5**), resulting in longer duration of detectable viral load (**Figure 6**). In pre-Omicron infections, the clearance period among unvaccinated individuals aged ≥65 years was 22.5 days (95% confidence interval: 21.2–23.8), compared with 17.5 days (17.3–17.7) in individuals aged <65 years. A similar trend was observed for Omicron infections, with clearance of 21.1 days (19.7–22.6) in unvaccinated individuals aged ≥65 years versus 15.9 days (15.7–16.2) in individuals aged <65 years. In contrast, the effect of age on peak viral load was moderate, with lower Ct values (reflecting higher viral load) in individuals aged ≥65 years, e.g. 16.0 (15.6–16.5) versus 17.6 (17.5–17.7) for Omicron unvaccinated ≥65 years versus <65 years, respectively.

**Figure 5:**
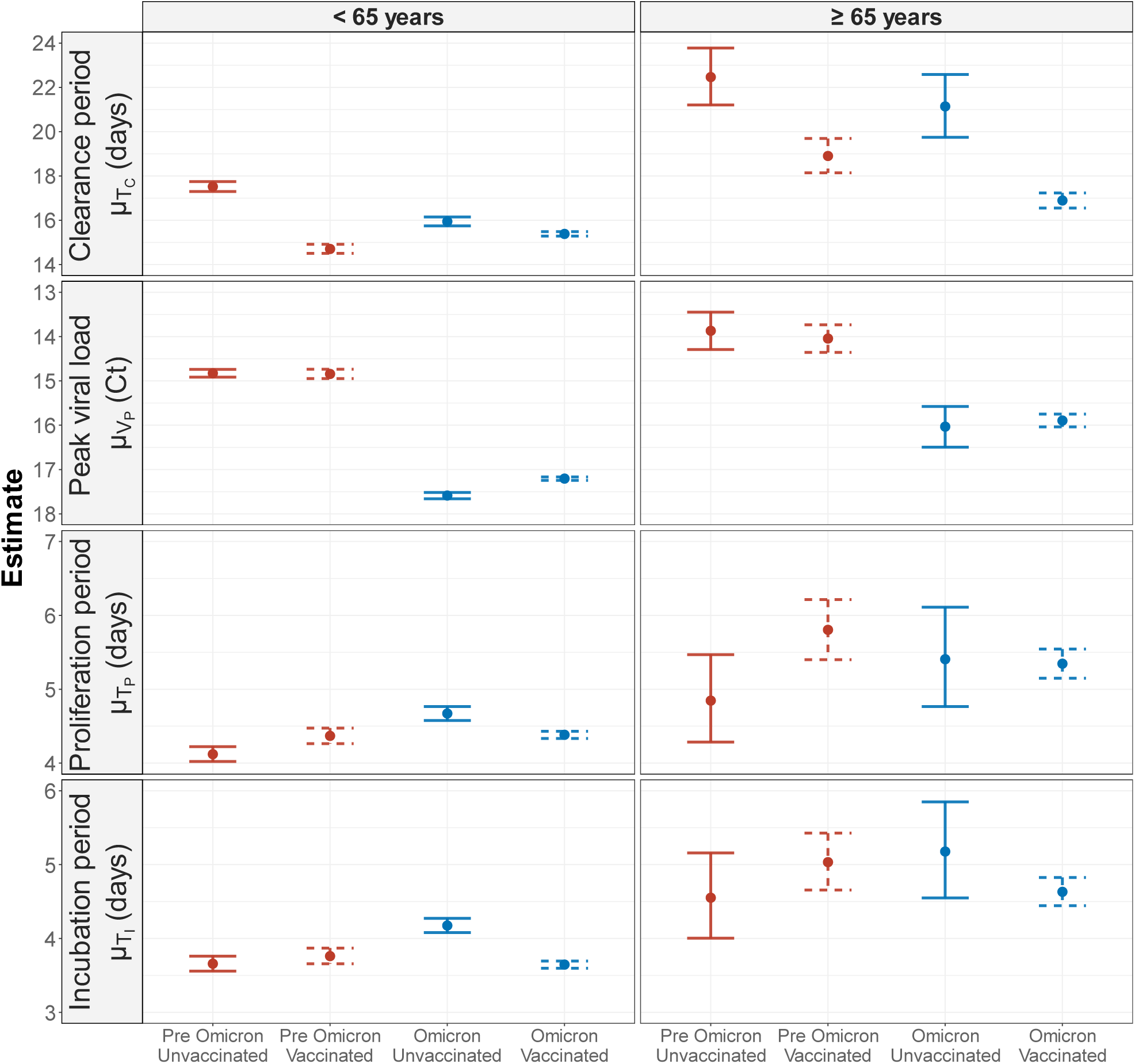
Viral dynamics parameters inferred from the data according to age, vaccination status and variant of infection. Estimated parameters of the viral dynamic model in the different population subgroups. Dots show the mean value, and vertical lines indicate the 95% confidence interval. Red: Pre-Omicron variants; blue: Omicron variants. Solid line: unvaccinated; dashed lines: vaccinated.

**Figure 6:**
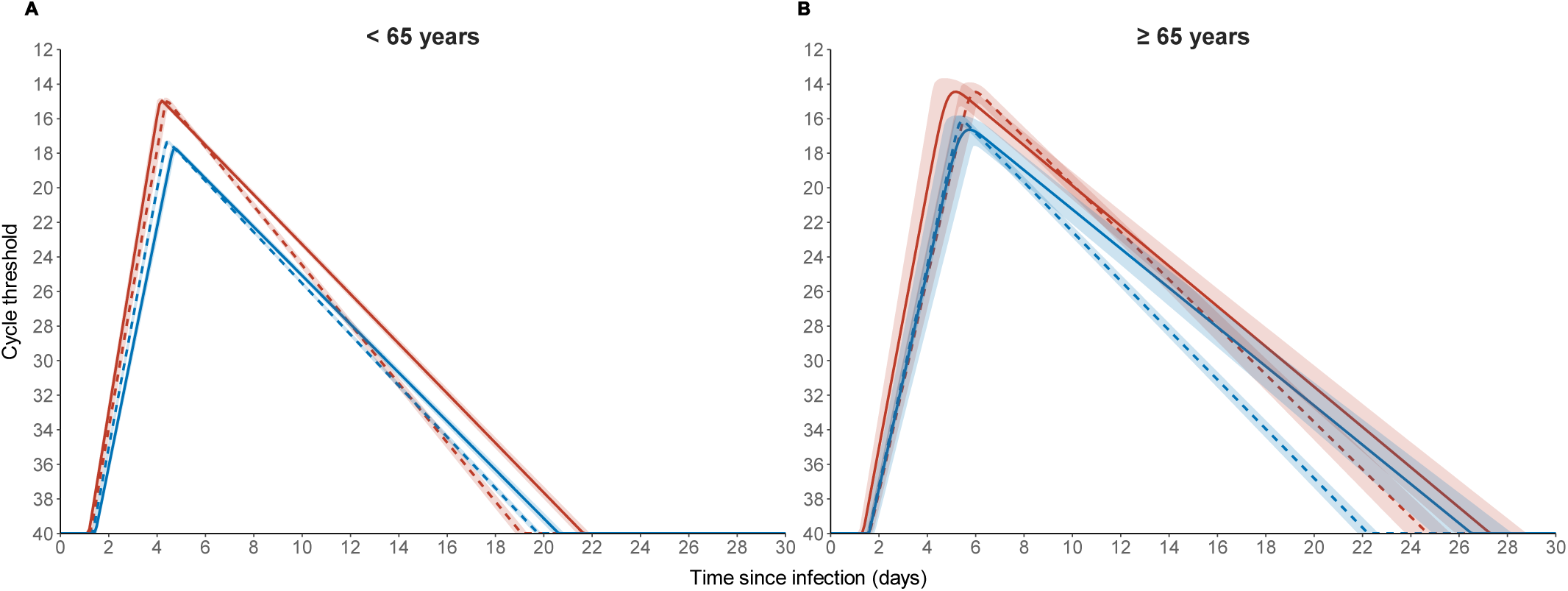
Predicted viral dynamics according to age, vaccination status and variant of infection. Model-based kinetics of the viral load value (in Ct) in the different population subgroups. Lines show the mean predicted Ct trajectory, with shaded areas indicating the 95% confidence interval. Red: Pre-Omicron variants; blue: Omicron variants. Solid line: unvaccinated; dashed lines: vaccinated.

Vaccination consistently shortened clearance period across variants and age groups, by approximately 2 to 4 days (**Figure 5**), resulting in shorter duration of detectable viral load (**Figure 6**). In pre-Omicron infections, clearance time decreased from 22.5 days (21.2–23.8) to 18.9 days (18.1–19.7) in individuals aged ≥65 years, and from 17.5 days (17.3–17.7) to 14.7 days (14.5–14.9) in individuals aged <65 years. In Omicron infections, the effect of vaccination on clearance was weaker among individuals aged <65 years, with a duration of 15.9 days (15.7-16.2) for the unvaccinated, and 15.4 days (15.3–15.5) for the vaccinated. The strongest reduction was observed in individuals aged ≥65 years, with a difference of 4.2 days between unvaccinated, 21.1 days (19.7-22.6), and vaccinated, 16.9 days (16.6-17.2). Vaccination had minimal impact on peak viral load, with differences consistently <0.5 cycle threshold.

Infection with an Omicron variant consistently decreased the peak viral load across vaccination and age groups, by approximately 2 to 3 Ct (**Figure 5**) compared to pre-Omicron variants. Pre-Omicron infections showed lower Ct values (higher viral loads) around 14–15, e.g. 14.8 (14.7–14.9) in unvaccinated individuals aged <65 years, compared with higher Ct values (lower viral loads) of 17–18 in Omicron, e.g. 17.6 (17.5–17.7) in unvaccinated individuals aged <65 years. Omicron infections tended to be associated with faster clearance periods than pre-Omicron infections, with the largest difference observed among vaccinated individuals aged ≥65 years, with a duration of 18.9 days (18.1- 19.7) in pre-Omicron infections, and 16.9 days (16.6-17.2) in Omicron infections.

## Discussion

Using a piecewise linear mixed-effects model on simulated data, key patterns of viral kinetics can be estimated from community-based testing data with good accuracy and precision. This modelling framework was subsequently applied to a very large dataset of PCR results collected in French community laboratories between July 2021 and March 2022, in order to investigate the effects of age, vaccination status, and variant of infection on viral load trajectories among symptomatic individuals. Our analyses showed that age >65 years was the primary factor affecting viral kinetic parameters, with an extended duration of the infection about 2 to 6 days. In addition, vaccination consistently shortened the clearance period across variants and age groups, by 2 to 4 days, without altering peak viral load. Both results suggest strong host immunity accelerates viral clearance but does not reduce peak viral load. In contrast, infections with the Omicron variants consistently exhibited lower peak viral load across vaccination and age groups, by approximately 2 to 3 Ct compared to Pre-Omicron variants. Furthermore, Omicron infections were cleared faster than pre-Omicron infections, with a maximum reduction of 2 days.

The factors identified here are consistent with findings from previous studies (8,9,25–27). Although several studies have used community-based data to model viral dynamics and identify factors shaping these trajectories(11,12,28), our study is the first to account for time since symptom onset to date infections and reconstruct viral kinetics. Moreover, it is the first, using community-based data, to robustly characterize the independent impact of Omicron and vaccination on within-host viral dynamics. Our work demonstrates that, beyond their role in epidemic surveillance, routinely collected PCR testing data can be leveraged to uncover novel determinants of viral dynamics. These data were gathered systematically on a daily basis, providing a low-cost means of building extensive datasets. This study retrospectively highlights the value of community testing data despite their inherent limitations, and supports their use in future outbreaks, especially with the growing availability of triplex PCR assays enabling the detection of multiple viruses from a single sample(29).

Our simulation study highlights the challenges of modelling large-scale datasets from community laboratories, where most data are available around symptom onset and longitudinal follow-up is limited. While modelling the entire population (i.e. including also the individuals with only negative PCR tests) in a Bayesian inference framework provided unbiased estimates across all parameters, as illustrated in our third scenario, this approach was computationally cumbersome (**Figure S6**) and was not feasible with our very large dataset. We therefore adopted a pragmatic strategy, focusing on modelling individuals with confirmed infections using frequentist inference in Monolix. This choice, however, comes at the cost of reduced accuracy and precision for early viral kinetic parameters. Nevertheless, the resulting estimation biases remained modest (approximately one day for early parameters and half a day for clearance, **Figure S4**) when considered in relation to the effects of covariate.

Another challenge, not addressed in our simulation study, relates to the source of the data, particularly clinical information such as symptoms, their onset time, or individuals’ vaccination status, which are often unreported or inconsistently recorded across tests. Even when available, these data are typically self-reported, leading to substantial amounts of missing data, and inconsistent information (**Figure S7**). Here we focused on individuals with information on age and sex, time of symptoms, vaccination, and variant, at the cost of excluding a large proportion of the dataset and not accounting for potential error in self-reported information. In particular, the timing of symptom onset was considered as perfectly known, even though this information was communicated as intervals of varying duration. We did not account for this uncertainty and instead imputed the median of the reported interval as the day of symptom onset. This simplification was mainly motivated by the fact that, in most cases, the uncertainty induced by the interval, was within one day. Nonetheless, a Bayesian inference framework could naturally incorporate this uncertainty in future work.

In our modelling, the testing behaviour was not investigated, which may limit the generalizability of the results due to potential selection bias. However, all data were collected during a period when the vast majority of PCR tests performed in France were fully reimbursed by the national health insurance(30), thereby limiting financial barriers to testing. Another factor limiting the generalization of our findings to the overall population is that the analysis focused exclusively on symptomatic infections, in order to date the infection. By restricting the modelling to these individuals, we excluded a substantial proportion of the dataset, as symptomatic cases represented only 45% of all positive tests, possibly leading to a reduction of precision in the proliferation period estimation. Among the 55% of infections without any declared symptoms, 33% corresponded to individuals who consistently reported being asymptomatic across all their tests, and 22% to those with unknown or missing symptom status at every test. The proportion of asymptomatic cases is consistent with prevalence estimates of around 40% reported in other studies(31). Furthermore, the Ct distribution of symptomatic individuals is shifted toward lower values (**Figure S8A**) when compared to asymptomatic individuals (**Figure S8B**), indicating potential differences in their underlying viral kinetics or the timing of their test.

We used a piecewise nonlinear model to reconstruct Ct trajectories. Mechanistic models in contrast may capture more complex dynamical phenomena such as viral load rebounds(32), and provide a better understanding of the biological impact of factors such as viral variants or vaccination on viral dynamics. However, the limited longitudinal follow-up per individual prevented the use of mechanistic models.

Finally, our study contributes to the broader effort of expanding methodological tools for analysing viral dynamics during epidemics. Such community-based laboratory data are increasingly available for several acute respiratory infections, particularly with the widespread adoption of multiplex PCR assays, which allow the systematic detection of multiple viruses from a single sample. Because these data are collected routinely and continuously, they represent a rich source of information at low cost. Leveraging them with appropriate statistical tools could help identify vulnerable subpopulations, monitor the impact of a new variant, or evaluate vaccine effectiveness in a population broadly representative of the general population. To support the wider use of community laboratory datasets, we recommend establishing clear frameworks for data collection, data sharing with research teams, and privacy protection.

## Acknowledgments

We thank Bob Carpenter and Charles Margossian (Flatarion Institute) for valuable discussions on modelling large datasets in Stan and on accounting for infection status in a Bayesian framework.

